# Unicellular and Multicellular Modes of Selection Impose Distinct Constraints on Cellular Phenotype Evolution

**DOI:** 10.64898/2026.09.16.751889

**Authors:** Mark Kim, Matt Pennell

## Abstract

Single-cell sequencing data have revealed that cellular phenotypes, such as gene expression states, are often low-dimensional, suggesting that cellular variation may arise from combinations of a smaller set of gene expression programs. A genome therefore defines a repertoire of cellular phenotypes that can be configured through different combinations of programs. However, organisms vary in how much of this repertoire is exposed to selection. In unicellular organisms, different phenotypes are often expressed across environments or life-cycle stages, so selection in a given context acts primarily through the phenotype expressed there. In multicellular organisms, multiple phenotypes can coexist within an individual and contribute jointly to fitness. Here, we use a geometric model to ask how selection acting through cellular phenotypes separately or jointly constrains the ability of a shared genome to evolve and maintain differentiated phenotypes across multiple functional demands. We vary the number of functional demands and how many corresponding phenotypes contribute jointly to fitness. We find similar evolutionary outcomes when demands are weakly divergent. Under strongly divergent demands, however, selection on one phenotype at a time leads to reduced differentiation as demands accumulate, even when sufficient programs are available. As more phenotypes contribute jointly to fitness, differentiation and performance improve. When all phenotypes contribute jointly, differentiation is maintained until demands outnumber programs. Our results suggest that how cellular phenotypes are organized in time and space can impose distinct constraints on the evolution of differentiation from a shared genome.

## Introduction

Cellular phenotypes, such as gene expression states, show remarkable variability despite being produced from a common genome within a species. In multicellular organisms, diverse cellular phenotypes are expressed across distinct cell types that coexist within tissues and organs, each expressing a different combination of genes encoded by the genome. In unicellular organisms, analogous variation is often expressed within individual cells over time, as cells switch between cell states, sometimes referred to as temporal cell types, in response to environmental conditions [Brunet et al., 2021, Dayel et al., 2011, Sebé-Pedrós et al., 2013], life-cycle stages [Brunet and King, 2017, de Mendoza et al., 2015, Fairclough et al., 2013, Sebé-Pedrós et al., 2017, Suga and Ruiz-Trillo, 2013], and stress conditions [Love and Wagner, 2022]. Such observations have motivated the hypothesis that spatial cell differentiation evolved through the integration of cell states that were already temporally differentiated in unicellular ancestors, an idea supported by comparative studies of animals and their unicellular relatives [Brunet and King, 2017, Mikhailov et al., 2009, Sebé-Pedrós et al., 2017, Zakhvatkin, 1949]. A similar pattern has been observed independently in volvocine algae, where genes that are diurnally regulated in unicellular lineages are partitioned between the soma and gonidia of *Volvox carteri* [Matt and Umen, 2018].

While multiple functionally distinct cellular phenotypes are produced from the same genome in unicellular and multicellular organisms alike, the temporal and spatial organization of these phenotypes poses different problems for adaptation. In a unicellular lineage, selection in a given environment or life-cycle stage acts primarily through the cellular phenotype expressed in that context. Mutations favored in one environment are exposed to selection through the phenotype expressed in that environment, while their effects on other phenotypes or functions may remain hidden [Chen and Zhang, 2020]. For example, in yeast evolution experiments, mutations selected under glucose limitation affect only a small number of fitness-relevant phenotypic dimensions near their evolution condition, while different components of their pleiotropic effects become fitness-relevant in more divergent environments [Ghosh et al., 2026, Kinsler et al., 2020]. Moreover, theory predicts that infrequent exposure can weaken selection on conditionally expressed genes or on phenotypes in rarely encountered environments [Draghi, 2021, Van Dyken and Wade, 2010]. Temporal separation among cellular phenotypes can therefore affect adaptation, by shifting which pleiotropic effects of a mutation are exposed to selection in a given context and by weakening selection on phenotypes that contribute to fitness only intermittently.

Multicellularity changes this relationship between selection and the cellular phenotypes that the genome can express. Multiple cellular phenotypes encoded by the genome can coexist within one individual, and therefore mutations in the genome can affect organismal fitness through several cell-level traits and functions at once. Effects on other cellular phenotypes that would remain hidden when those phenotypes are expressed in different contexts can therefore become fitness-relevant when several phenotypes contribute to the fitness of the same individual. On the one hand, multicellularity may enable selection to maintain distinct cellular phenotypes by compartmentalizing poorly compatible functions among cells and stabilizing a division of labor that is inaccessible to unicellular organisms [Arendt et al., 2009, Ispolatov et al., 2012, Rueffler et al., 2012]. At the same time, multicellularity can impose additional constraints, as one genome must support multiple functional demands simultaneously. As such, adaptation can become more difficult due to the increasing cost of pleiotropy as mutations affect more phenotypic characters under selection [Orr, 2000, Wagner et al., 2008, Wagner and Zhang, 2011, Wang et al., 2010, Welch and Waxman, 2003]. While studies of selection on multiple traits have shown that evolutionary responses can depend on how selection acts jointly across traits [Davidowitz et al., 2016, Zijlstra et al., 2003], few have directly compared how the order or simultaneity of selection on multiple functional demands shapes which evolutionary outcomes remain accessible in multicellular or unicellular eukaryotes. Nevertheless, evidence from other systems suggests that the distinction can be important. For instance, experiments in bacteriophage show that separate and joint selection on multiple phenotypes can make different outcomes of adaptive evolution accessible through different paths, with lineages first selected on growth alone and subsequently on both growth and capsid stability reaching states that were less accessible when both traits were selected simultaneously from the beginning [Sackman and Rokyta, 2019].

Hence, differences in how cellular phenotypes are organized can impose different constraints on adaptation from a shared genome. When cellular phenotypes contribute to fitness in different environmental or life-cycle contexts, effects of a mutation on phenotypes that are not currently contributing to fitness can remain hidden from selection. When multiple cellular phenotypes contribute jointly to fitness, the effects of the same mutation across those phenotypes are evaluated together, potentially imposing additional pleiotropic constraints [Chebib and Guillaume, 2022, Espinosa-Soto and Wagner, 2010]. This raises two related questions. How can a single genome evolve and maintain differentiated cellular phenotypes? And how does this capacity depend on the number of environmental (in unicellular organisms) or functional (in multicellular organisms) demands it must satisfy and whether the corresponding phenotypes contribute to fitness separately or jointly?

Addressing these questions requires a model in which a single genome can generate multiple cellular phenotypes from a shared set of functional components. Recent single-cell transcriptomic studies show that variation in gene expression across cell types and cellular states often has such a modular structure, with genes co-expressed in recurring sets termed gene expression programs (GEPs), each associated with a particular cellular function or activity [Kotliar et al., 2019, Musser et al., 2021, Parker and Pennell, 2025, Sebé-Pedrós et al., 2018, Tarashansky et al., 2021, Trapnell, 2015, Wagner et al., 2016]. Models of cell-type evolution have similarly described cellular phenotypes as combinations of functional modules that can be reused or partitioned among cellular contexts [Achim and Arendt, 2014, Arendt et al., 2019, 2016].

To this end, we build on the toolbox model of Tikhonov et al. [2020]. In this model, a genome encodes a finite set of reusable components that can be combined in different proportions to generate phenotypes adapted to different environmental demands. We extend the original formulation from alternation between two environments to an arbitrary number of environmental or functional demands and vary how many corresponding phenotypes contribute to fitness at the same time. Using this model, we simulate adaptive evolution and ask how the number of demands placed on a shared genome and the extent to which those demands contribute jointly to fitness constrain the evolution of differentiated cellular phenotypes and their performance.

### Genotype, regulation, and phenotype

We represent the genome as a collection of *L* genetic loci whose effects on each of *K* programs are encoded by a binary matrix, *G ∈ {*0, 1*}L×K* . The entry *G_ℓk_*indicates whether the allelic state at locus *ℓ* affects phenotype when program *k* is deployed. In our model, programs represent intermediate phenotypic components analogous to the GEPs described in single-cell biology [Kotliar et al., 2019, Parker and Pennell, 2025]. For each environmental or functional demand, a task-specific cellular phenotype is generated by combining these programs, **z** = *G***a**, where program usage is denoted by the non-negative vector **a** *≥* 0. This mapping is illustrated in Figure 1A.

**Figure 1:**
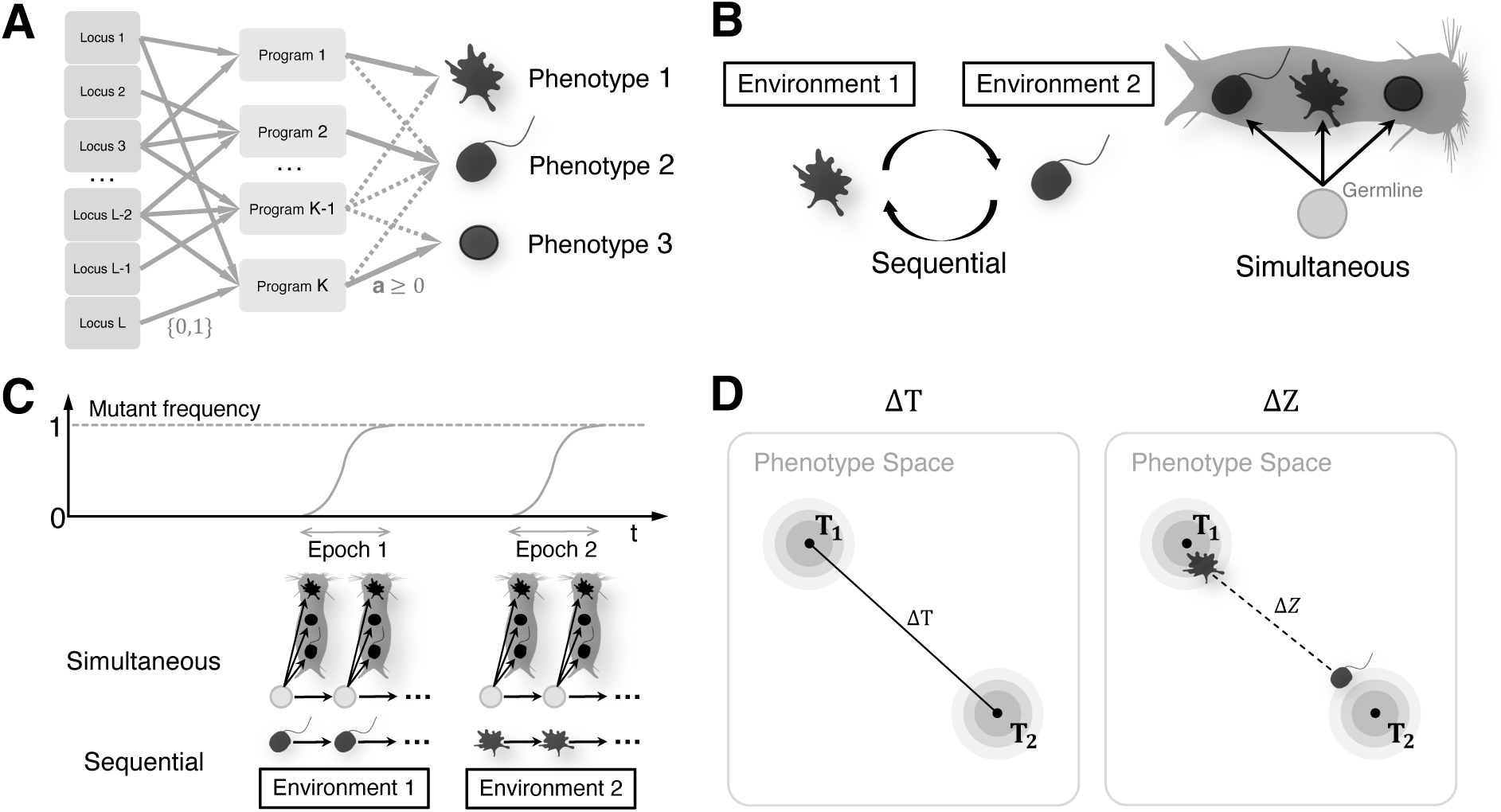
Schematic of the model and selection regimes. **A.** Genotype-to-phenotype map. A genome with *L* loci encodes *K* programs, which are combined with non-negative weights **a** to produce a task-specific phenotype. **B.** Sequential and simultaneous selection. Fitness depends on performance on one task at a time under sequential selection and on all tasks jointly under simultaneous selection. **C.** Evolutionary dynamics in the strong-selection weak-mutation (SSWM) regime. Mutations fix successively. Under sequential selection, one task is drawn during each selective epoch; under simultaneous selection, all tasks contribute to fitness during every epoch. **D.** Phenotypic differentiation. **T**_1_ and **T**_2_ are task optima separated by Δ*T*. The corresponding phenotypes are separated by Δ*Z*. The ratio Δ*Z/*Δ*T* measures differentiation relative to the separation between task optima. **Alt text:** Four panels illustrate the model and selection regimes. Panel A shows genetic loci connecting to programs, which combine with non-negative weights to produce three cellular phenotypes. Panel B contrasts a single cell alternating between two environments under sequential selection with a multicellular organism containing three coexisting differentiated cells under simultaneous selection. Panel C shows mutant frequency rising from zero to one through successive fixations. Below, the simultaneous regime shows the organism in both selective epochs, whereas the sequential regime shows different cellular phenotypes in different environments. Panel D compares the separation between two task optima, labeled Δ*T*, with the smaller separation between the corresponding evolved phenotypes, labeled Δ*Z*.

We define a set of phenotypic optima as *L*-dimensional non-negative target vectors, **T***_j_ ∈* R*^L^*+. Each optimum represents the expression profile required to perform task *j* and is normalized to unit length, so distances among optima depend on relative expression patterns rather than overall expression magnitude. We use the term task throughout, but the same formalism also applies when **T***_j_* is more naturally interpreted as the optimum imposed by environment *j*.

For task *j*, the genome can deploy the phenotype that optimally approximates the phenotypic optimum for that task. This assumes that expression is sufficiently plastic to always realize the optimal combination of programs for whichever task is active. Because **a***∗* minimizes the residual for each task by construction, the performance we report is an upper bound on what a given genome and program repertoire can achieve. We denote the corresponding coefficient vector by **a***∗*, chosen to minimize

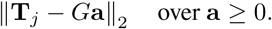

The residual *d_j_*(*G*) = ‖T*_j_* – *G***a***_j_*\*‖ measures how well the genome’s program repertoire approximates the task demand, and task performance is

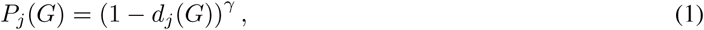

where *γ >* 0 parametrizes how performance declines with mismatch [Adler et al., 2019]. We focus on linear decay (*γ* = 1) for the main results and examine the effects of relaxing this assumption in the robustness analyses (Figure S3).

### Selection regimes

The number of tasks contributing jointly to fitness defines the evolutionary regimes we vary (Figure 1B). We use the term selective epoch to denote an interval during which a fixed set of task-specific phenotypes contributes to fitness, thereby determining which phenotypic effects of new mutations are exposed to selection. During each selective epoch, a subset *S ⊆ {*1*, . . ., T }* of size *m* is drawn uniformly at random from all subsets of size *m*, corresponding to the task-specific phenotypes that contribute to fitness during that epoch. When *m* = 1, fitness depends on a single active task during each selective epoch. We refer to this as the *sequential* regime, recovering the sequential fluctuating-environment setup described in Tikhonov et al. [2020]. When *m* = *T*, all tasks contribute jointly to fitness during every selective epoch, which we term the *simultaneous* regime. The intermediate values 1 *< m < T* interpolate between these theoretical limits. Biologically, the simultaneous limit is motivated by multicellular organisms in which differentiated cell types coexist within the same individual, while the sequential limit is a limiting case that isolates the consequence of only one phenotype contributing to fitness during each selective epoch.

Fitness is defined as a power mean of task performance on *S*,

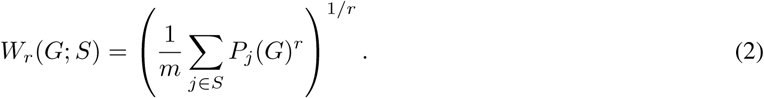

Inactive tasks (*j ∈/ S*) do not contribute to fitness during the current selective epoch. The geometric mean is recovered in the limit *r →* 0,

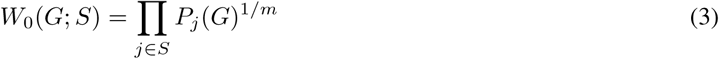

which we use for the main results as a simple multiplicative aggregation of performance across active tasks. Other forms of fitness aggregation may be more appropriate for other biological contexts, so we also examine a negative power mean (*r* = *−*2), which increases the influence of the poorest-performing active task, in the supplementary analysis (Figure S2).

### Evolutionary dynamics and simulation

We simulate evolutionary dynamics in the strong-selection weak-mutation (SSWM) regime, where the population is effectively monomorphic and adaptation proceeds by the sequential fixation of new mutations [Gillespie, 1983, McCandlish and Stoltzfus, 2014], as illustrated in Figure 1C. We initialize the genome by independently drawing each entry from a Bernoulli distribution, *G_ℓk_ ∼* Bernoulli(*ρ*), with *ρ* = 0.25. For the main analyses, *K* = 4, so each locus affects one program on average. We keep the same initial density when varying the number of programs, and the effect of a denser initialization (*ρ* = 0.5) on our main results is explored in the supplementary analysis (Figure S5). Each mutation changes one entry of the genotype matrix from 1 to 0 or from 0 to 1, changing whether a locus affects a particular program. We do not assign a unique molecular interpretation to this mutation operation, since the same change in the genotype–phenotype map could arise through several underlying mechanisms. For instance, it could represent a coding mutation whose effect is specific to the cellular context in which the program is deployed, a cis-regulatory mutation that alters the locus’s contribution to that program, or a trans-acting mutation elsewhere in the genome that modifies the locus’s program-specific effect through another gene or regulatory factor.

For each mutant genotype, the optimal deployment coefficients **a***∗* are recalculated for each task. We consider only beneficial mutations, which fix with probability

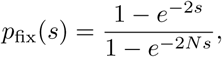

where *N* is the effective population size of haploid individuals [Gillespie, 2004]. The selection coefficient *s* of the mutant genome *G^′^* relative to the currently fixed genome *G* under the active task set *S* is

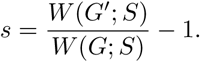

Finally, for a given active task set, the substitution process associated with each beneficial mutation *i ∈ B* is modeled as an independent Poisson process with rate *Nµp*_fix_(*s_i_*). The waiting time to the next substitution is drawn from an exponential distribution

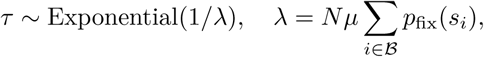

and the substituting mutation is sampled with probability

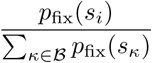

[Gillespie, 1977]. If no beneficial mutation is available under the current active task set, a new active task set is drawn and the mutation-selection step is repeated.

For all our simulation results, we use *N* = 104 and the per-site per-generation mutation rate *µ* = 10*−*7. Because initial genomes are generated independently of the task optima, populations begin without prior adaptation to the imposed demands. We compare populations after 50 and 400 substitutions to characterize earlier and later stages of adaptation, respectively. The choice of comparison points is described in Supplementary Section 1.5, and simulation stopping criteria and the treatment of populations that stopped earlier are described in Supplementary Section 1.2 (Figure S1A–C).

### Task ensemble construction

We use a symmetric Dirichlet distribution to generate a set of non-negative vectors that define the phenotypic optima for fitness-relevant tasks. For each task ensemble of size *T*, target profiles *{***T***_j_}T* are drawn independently from

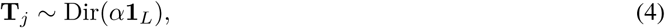

and subsequently normalized to unit length. The concentration parameter *α* controls the sparsity of the target vectors, with *α <* 1 producing sparse, peaked vectors and *α >* 1 producing dense, near-uniform vectors. To vary the mean pairwise target separation Δ̄*T̄*, defined as

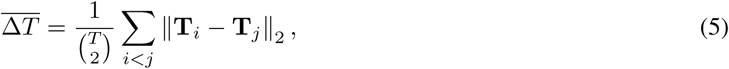

the parameter *α* is numerically calibrated for each prescribed value of Δ*T* .

### Simulation design and measures of phenotypic evolution

For the main analyses, we varied the number of task-specific phenotypes contributing jointly to fitness (*m*), the number of tasks (*T*), and the mean pairwise target separation (Δ*T*), while keeping the dimensions of the genotype-to-phenotype map fixed at *L* = 100 loci and *K* = 4 programs (see Figure S4 for supplementary analyses with *K* = 6 and *K* = 8). Parameter values for the main and supplementary analyses are summarized in Supplementary Table S1.

Each replicate population was assigned an independently drawn task ensemble and initial genotype matrix, and for each task ensemble size (*T*), the same replicate was evolved under every value of the simultaneity parameter, *m ∈ {*1*, . . ., T }*. We characterized the evolved phenotype repertoire under multiple functional demands using two metrics. *(i)* Phenotypic differentiation measures the extent to which differences among task-specific phenotypes track differences among the corresponding task demands, while *(ii)* optimization measures how closely those phenotypes match their corresponding task optima.

To quantify phenotypic differentiation, we associated each task *j* with the phenotype **z***_j_*(*G*) = *G***a***∗* that optimally approximates the corresponding task optimum **T***_j_* (see section Genotype, regulation, and phenotype). We quantified the mean pairwise distance among these evolved phenotypes,

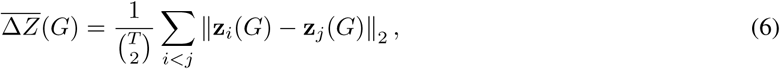

and normalized this quantity by the mean pairwise separation among that replicate’s own task optima, Δ̄*T̄*, to obtain the degree of differentiation Δ̄Z*̄*/Δ̄*T̄*. This quantity is bounded between zero and one, 0 ≤ Δ̄Z*̄*/Δ̄*T̄* ≤ 1 (see Supplementary Section 1.4). A value of one indicates that the evolved phenotypes are separated to the same extent as the task optima, whereas values below one indicate that the genome deploys more similar phenotypes across tasks than the task demands themselves would require, reflecting incomplete differentiation (Figure 1D). We quantified optimization as the extent to which task demands were met in absolute terms. Combining the per-task residuals into a vector **d**(*G*) = (*d*_1_(*G*)*, . . ., d_T_* (*G*)), we defined the degree of optimization

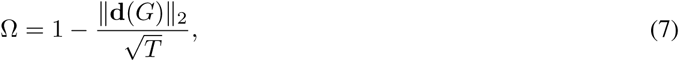

where √*T* is the maximum attainable norm of **d**(*G*). Thus, 0 *≤* Ω *≤* 1 (see Supplementary Section 1.4), with Ω = 1 indicating exact matching of every task optimum and lower values indicating greater residual mismatch across tasks.

## Results and Discussion

### Sequential selection limits differentiation as tasks accumulate

We first consider the sequential regime (*m* = 1), which extends the alternating-environment setup of Tikhonov et al. [2020] from two environmental demands to an arbitrary number of environmental or functional demands. In our model, *m* = 1 represents the theoretical limit in which only one task-specific phenotype contributes to fitness during each selective epoch. To instantiate this limit biologically, we consider cases in which the fitness contributions of different cellular phenotypes are separated across the timescale over which mutations experience selection (Figure 2A). For example, several dinoflagellate taxa form dormant resting cysts that can persist in marine sediments for decades before germinating, providing an extreme case of temporal separation between cell states [Lundholm et al., 2011]. Since only one of *T* task-specific phenotypes contributes during any given selective epoch, each phenotype contributes in an expected fraction 1*/T* of epochs. As *T* increases, *m* = 1 therefore represents an increasingly restrictive case in which a growing phenotypic repertoire is evaluated one phenotype at a time.

**Figure 2:**
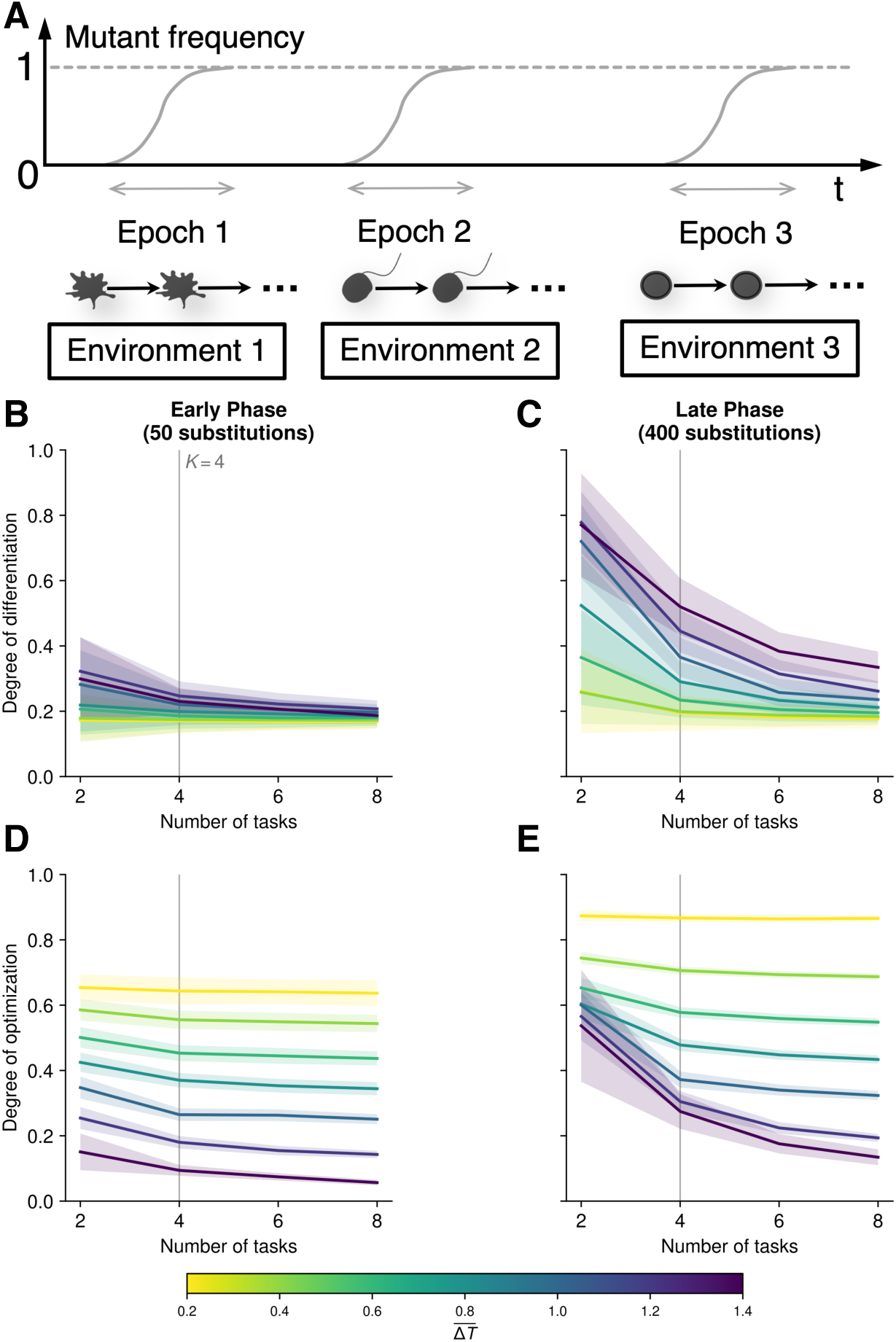
Under sequential selection, differentiation declines before tasks outnumber programs. **A.** Sequential selection (*m* = 1). One task-specific phenotype contributes to fitness during each selective epoch. **B, C.** Phenotypic differentiation, Δ̄Z*̄*/Δ̄*T̄*, after 50 (**B**) and 400 (**C**) substitutions. Δ̄Z*̄* is the mean pairwise distance between evolved phenotypes, and Δ̄*T̄* is the mean pairwise distance between task optima. A value of *√*one indicates that phenotypes are as separated as their task optima. **D, E.** Optimization index, Ω = 1 – ‖*d*(*G*)‖_2_/√*T*, after 50 (**D**) and 400 (**E**) substitutions. Colors indicate target separation, Δ̄*T̄*. Vertical lines mark *T* = *K* = 4. Lines show means and shaded regions indicate ±1 standard deviation across 200 replicate populations, each with an independently drawn task ensemble and initial genotype. *L* = 100, *K* = 4, *γ* = 1, *r* = 0, *ρ* = 0.25, *µ* = 10*−*7, and *N* = 104. **Alt text:** A schematic of sequential selection sits above four plots comparing 50 and 400 substitutions. The schematic shows successive selective epochs, each with a different environment and a different task-specific phenotype contributing to fitness. Differentiation and optimization are plotted against task number, with colors running from yellow for weakly separated optima to purple for strongly separated optima. Grey vertical lines mark four tasks, equal to the number of programs. Differentiation is low at 50 substitutions. Later, strongly separated optima produce greater differentiation at two tasks, but differentiation and optimization decline as tasks accumulate. Weakly separated optima retain low differentiation and high optimization. Shading shows one standard deviation across replicate populations.

In the early stage of adaptation, mean phenotypic differentiation was low under every condition examined (Figure 2B), while optimization already depended strongly on target separation (Figure 2D). When the task optima were close together, the degree of optimization was relatively high and changed little as tasks accumulated. As the task optima became more strongly separated, performance declined substantially. Optimization also declined as task number increased, particularly when task optima were well separated (Figure 2D). Thus, early in evolution, task divergence primarily affected how closely the deployed phenotypes approached their optima, while substantial differentiation had not yet evolved under any of the conditions examined.

In the later stage of adaptation, the observed patterns differed qualitatively depending on the separation among task optima (Figure 2C,E). When the task optima were close together (e.g., Δ̄*T̄* = 0.2), phenotypic optimization increased substantially between the early and late stages (Figure 2D–E, yellow line). Differentiation declined only slightly with task number and remained low in absolute terms (Figure 2B–C, yellow line). This result shows that in our simulations, the genome could improve its fit to several closely related optima while maintaining relatively similar phenotypic states across tasks. When the task optima were strongly separated, however, a qualitatively different pattern emerged (see Δ*T* = 1.4, for instance). With a small number of tasks, both differentiation and phenotypic optimization increased substantially from the early to the late stage of adaptation (Figure 2B–E). The magnitude of this improvement became progressively smaller as the number of tasks increased, and late-stage differentiation and optimization declined with task number (Figure 2C,E, purple line). Thus, when the demands were strongly divergent, a genome evolved under sequential selection produced well-differentiated and well-optimized phenotypes when the number of tasks was small, but differentiation and optimization deteriorated as the number of demands requiring distinct task-specific phenotypes increased.

Interestingly, differentiation declined even when tasks did not outnumber programs (*T ≤ K*; Figure 2C; see also Figure S4). The finite number of programs therefore cannot by itself explain the loss of differentiation under sequential selection. Because *m* = 1, only one task-specific phenotype contributes to fitness during each selective epoch (Figure 2A), so any particular phenotype contributes in an expected fraction 1*/T* of selective epochs. A mutation favored through the currently active phenotype can also alter other task-specific phenotypes through pleiotropy, but those effects do not influence the fate of the mutation when the corresponding phenotypes are inactive [Bakerlee et al., 2021, Kinsler et al., 2020]. As *T* increases, each phenotype contributes to fitness less frequently, so a larger fraction of a mutation’s effects across the phenotype repertoire is hidden from selection during the selective epoch in which that mutation arises and fixes. Mutations can therefore be favored for their effects on the active phenotype even when their effects on other phenotypes reduce the genome’s ability to maintain differentiation across tasks. This parallels the weakened selection expected under conditional gene expression [Van Dyken and Wade, 2010], extended here from a single conditionally expressed gene to a repertoire of phenotypes that share a genetic architecture.

This result shows that limited exposure of the phenotype repertoire to selection can constrain differentiation even when the genome has a sufficient number of available programs to generate distinct phenotypes for the different demands. For unicellular lineages with multiple cell states, these results suggest that the availability of programs for distinct functions may not be sufficient to maintain differentiation when selection acts on the corresponding phenotypes separately. We revisit this point below, where we simulate 1 *< m < T* directly.

### Simultaneous selection preserves differentiation until tasks outnumber programs

We next consider *m* = *T*, the opposite theoretical limit, in which all task-specific phenotypes contribute jointly to fitness during every selective epoch (Figure 3A). Spatially differentiated cell types in multicellular organisms provide a clear biological motivation for this regime, yet spatial coexistence is not a necessary condition. Temporally expressed cellular phenotypes can also approach or reach the *m* = *T* limit when all of them recur within the timescale over which new mutations experience selection, so that their contributions to fitness are jointly relevant to the fate of a mutation. The obligate asexual cycle of *Plasmodium* provides such an example, as parasite cells repeatedly pass through ring, trophozoite, and schizont states [Howick et al., 2019]. A mutation inherited across successive generations is therefore repeatedly exposed to selection through all three states, so that its fitness effects across those states jointly influence its evolutionary fate. Across successive stages of a temporally structured life cycle, fitness can accumulate multiplicatively over time [Jasmin and Kassen, 2007], which we represent in our simulations using the geometric mean to combine performance across task-specific phenotypes that contribute within the same selective epoch (see Selection regimes). Spatial differentiation provides another instantiation of this regime, as in *Volvox carteri*, where somatic cells and germline gonidia differentiate during embryogenesis and coexist within the same organism for the remainder of the generation [Matt and Umen, 2016]. With both cell types present throughout the generation, mutations can affect organismal fitness through both cell types within the same selective epoch.

**Figure 3:**
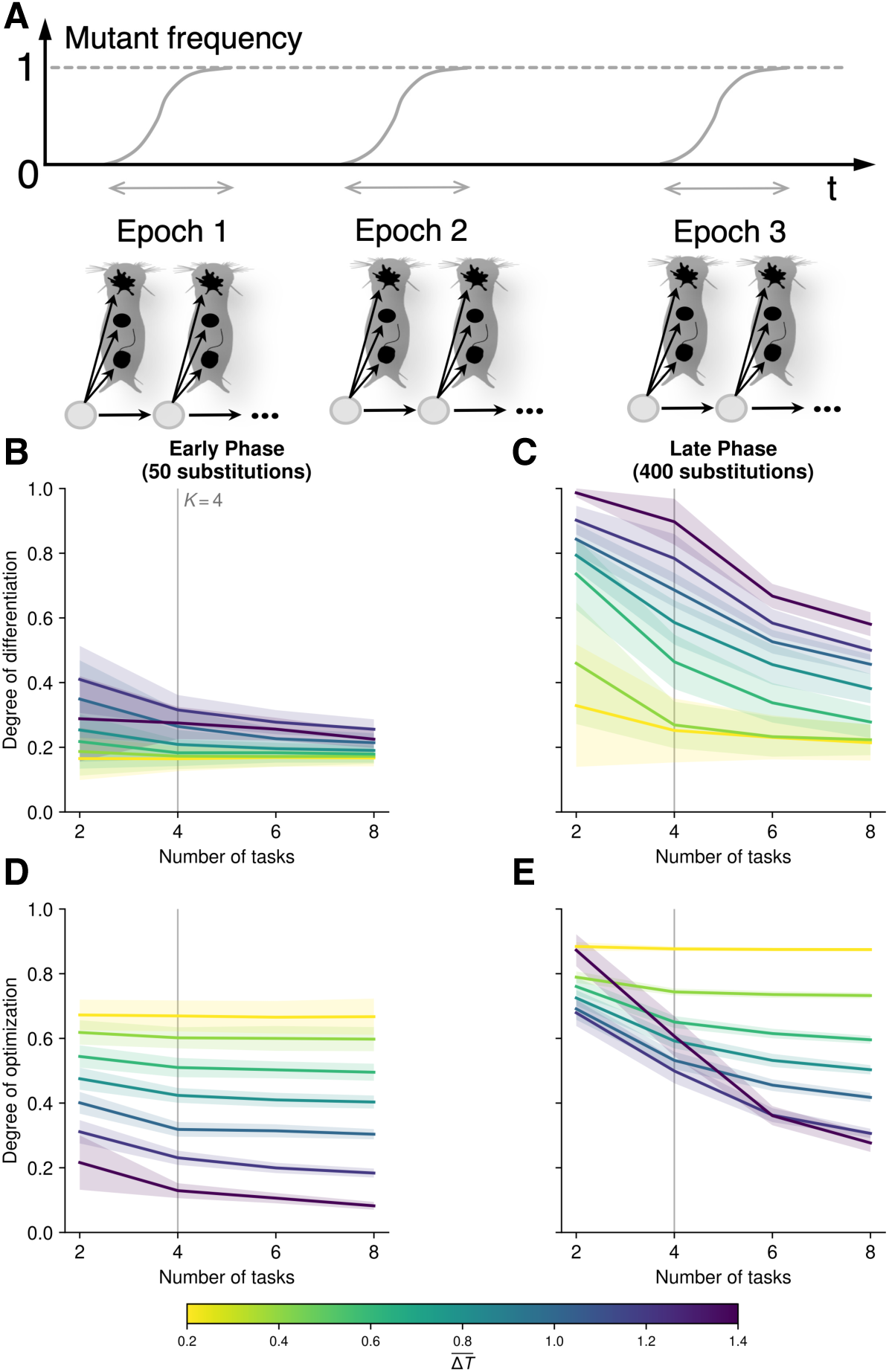
Under simultaneous selection, differentiation is maintained until tasks outnumber programs. **A.** Simultaneous selection (*m* = *T*). All task-specific phenotypes contribute jointly to fitness during every selective epoch. **B, C.** Phenotypic differentiation, ΔZ*̄*/Δ*T̄*, after 50 (**B**) and 400 (**C**) substitutions. **D, E.** Optimization index, Ω, after 50 (**D**) and 400 (**E**) substitutions. Measures are defined in Figure 2. Colors indicate target separation, Δ̄*T̄*. Vertical lines mark *T* = *K* = 4. Lines show means and shaded regions indicate 1 standard deviation across 200 replicate populations. Parameters are as in Figure 2. **Alt text:** A schematic of simultaneous selection sits above four plots comparing 50 and 400 substitutions. The schematic shows successive selective epochs in which all task-specific phenotypes contribute to fitness together. Differentiation and optimization are plotted against task number, with colors running from yellow for weakly separated optima to purple for strongly separated optima. Grey vertical lines mark four tasks, equal to the number of programs. Differentiation is low at 50 substitutions. At 400 substitutions, strongly separated optima yield differentiation close to one when tasks do not outnumber the four programs, followed by a decline when tasks outnumber programs. Optimization also improves during adaptation but declines with task number. Weakly separated optima retain high optimization. Shading shows one standard deviation across replicate populations.

In the simulations, the early stage of adaptation under simultaneous selection resembled that under sequential selection. Phenotypic differentiation remained low (Figure 3B), while optimization declined as target separation increased (Figure 3D). By the later stage of adaptation, however, the two regimes had diverged substantially (Figures 2C,E and 3C,E). When *T ≤ K*, phenotypic differentiation remained high even for strongly separated task optima, approaching the degree of separation among the task demands themselves (Figure 3C; see the strongly separated task ensembles for *T ≤* 4). Phenotypic optimization also increased substantially over the evolutionary trajectory (Figure 3D,E), although within the late phase it declined progressively as the number of tasks increased (Figure 3E). Having at least as many programs as tasks therefore allowed simultaneous selection to maintain strongly differentiated task-specific phenotypes, even though those phenotypes became less optimized as the number of tasks increased. When tasks outnumbered programs, both differentiation and optimization declined, particularly when task optima were strongly separated (Figure 3C,E).

This contrast with sequential selection shows that whether program number acts as a threshold for maintaining differentiation depends on the selection regime (Figures 2C and 3C; Figure S4A,B). At *m* = 1, only one task-specific phenotype contributes to fitness during each selective epoch, so effects on the others remain cryptic until those phenotypes become active, and differentiation declines as tasks accumulate even when tasks do not outnumber programs. At *m* = *T*, every task-specific phenotype contributes to fitness at once, so effects that would otherwise remain cryptic become fitness-relevant immediately, and mutations are selected on their joint contribution across phenotypes. When effects across all phenotypes are exposed to selection, the number of available programs becomes the main constraint on maintaining divergent phenotypes. Phenotypic optimization nevertheless declined progressively as the number of tasks increased, even while differentiation remained high for *T ≤ K*.

It is important to note that how performance across active functions should be aggregated into fitness (equation 2) is itself a biological question. The geometric mean has a natural interpretation when performance across sequentially realized functions contributes multiplicatively to growth, as can occur when cell-level functions cycle through fast, repeated use within a generation. Organisms whose cell types depend on each other more tightly, so that failing any single function is disproportionately costly, might instead be better described by a minimum or soft-minimum function, which penalizes one weak function much more heavily than the geometric mean does. We tested whether our results depend on this choice by replacing the geometric mean with a negative power mean (*r* = *−*2), which weights the poorest-performing active task most heavily. Sequential selection still lost differentiation rapidly as tasks accumulated, while simultaneous selection continued to maintain substantially higher differentiation and optimization at strongly separated task optima (Figure S2), so the contrast between the two regimes does not depend on which of these aggregation rules is used.

Our finding was similarly robust to the curvature of the performance function. Theory on the evolution of division of labor suggests that the shape of the performance function can affect whether specialization is favored [Rueffler et al., 2012]. We therefore tested whether the contrast between selection regimes persists when performance declines more sharply with distance from a task optimum (equation 1). Sharpening the decline in performance with phenotypic mismatch (*γ* = 4) did preserve the pattern, with sequential selection losing differentiation rapidly as tasks accumulated while simultaneous selection maintained it (Figure S3).

### Differentiation can evolve without fully simultaneous selection

Between the sequential and simultaneous limits, more than one but fewer than all task-specific phenotypes can contribute jointly to fitness during the same selective epoch. We therefore asked whether the difference between *m* = 1 and *m* = *T* emerges only under full simultaneity or develops progressively as *m* increases. We find that intermediate values of *m* recover part of the differentiation and optimization achieved under simultaneous selection. When task optima were close together, half-simultaneity (*m* = *T/*2) recovered much of the gain in both differentiation and optimization across *T* = 4, 6, and 8 (Figure 4A,D). In contrast, when task optima were strongly separated, the same degree of simultaneity recovered a smaller fraction of either gain (Figure 4C,F). Thus, differentiation and optimization can improve before all phenotypes contribute jointly to fitness, but stronger divergence among functional demands requires broader joint contribution to achieve comparable gains.

**Figure 4:**
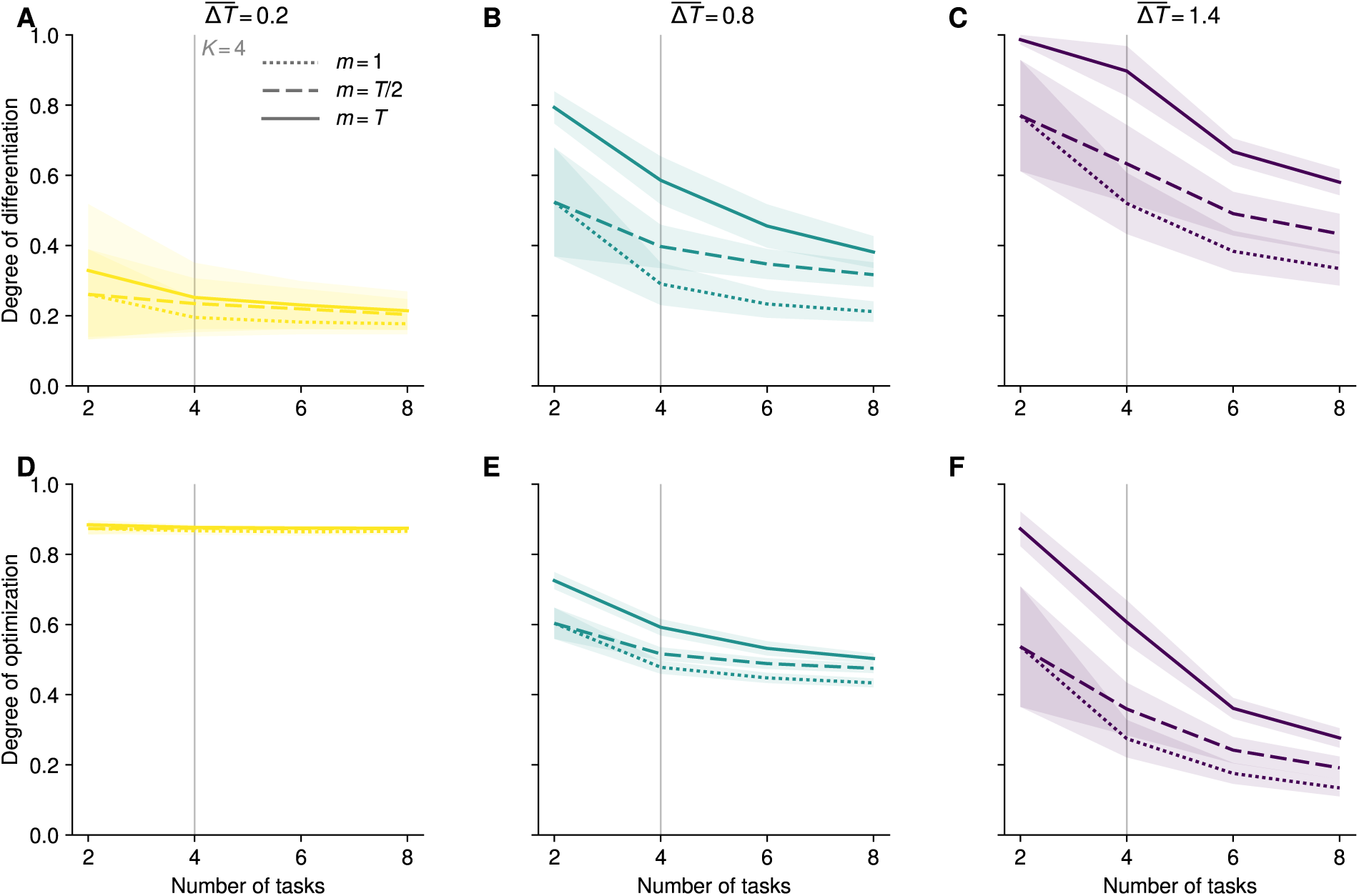
Joint selection on part of the phenotype repertoire improves differentiation and performance. **A–C.** Phenotypic differentiation, ΔZ*̄*/Δ*T̄*, after 400 substitutions. **D–F.** Optimization index, Ω, at the same stage. Columns and colors indicate target separation, Δ*T̄* = 0.2, 0.8, and 1.4. Dotted, dashed, and solid lines correspond to *m* = 1, *m* = *T/*2, and *m* = *T*, respectively. At *T* = 2, *m* = *T/*2 coincides with sequential selection. Vertical lines mark *T* = *K* = 4. Lines show means and shaded regions indicate ±1 standard deviation across 200 replicate populations. Measures and parameters are as in Figure 2. **Alt text:** Six plots show differentiation in the top row and optimization in the bottom row against task number. Columns represent weak, intermediate, and strong separation among task optima. Dotted, dashed, and solid curves represent selection on one task-specific phenotype, half of the task-specific phenotypes, and all task-specific phenotypes, respectively. Selection on half the tasks generally improves both measures over sequential selection, but recovers a smaller fraction of the full simultaneous advantage when optima are strongly separated. Grey vertical lines mark four tasks, equal to the number of programs. Shading shows one standard deviation across replicate populations.

Intermediate and fully simultaneous selection differed in how their advantages over sequential selection changed as tasks accumulated. When task optima were strongly separated, the advantage of *m* = *T* over sequential selection was greatest while *T ≤ K* and declined after the number of tasks exceeded the number of programs (Figure 4C,F; Figure S4). At *m* = *T/*2, in contrast, the corresponding advantage changed only modestly across *T* = 4, 6, and 8, with no comparable change around *T* = *K* (Figure 4C,F). These results suggest that the number of available programs becomes a stronger constraint on differentiation as more phenotypes contribute jointly to fitness. At intermediate *m*, selection still acts on only part of the phenotype repertoire during each epoch, and differentiation remains limited even when enough programs are available to represent the task demands.

We then examined gains in differentiation and optimization at every value of *m* (Figure S6). When task optima were strongly separated, both gains increased up to the simultaneous limit. At weaker separation, however, the largest gains occurred when slightly fewer than all phenotypes contributed jointly to fitness. Joint selection on part of the repertoire was therefore sufficient to improve both measures, and including every phenotype did not always improve them further. These comparisons were made after 400 substitutions, by which point populations under simultaneous selection had reached local fitness peaks, while many populations at intermediate *m* continued to change (see Supplementary Section 1.5). Differences in the course of adaptation may therefore contribute to the greater gains at intermediate *m* under weaker target separation.

These results suggest that the evolutionary consequences of temporal and spatial differentiation depend on how much of the phenotype repertoire contributes to fitness within the same selective epoch. Temporal separation among phenotypes will tend to reduce this joint contribution when different phenotypes recur across different selective epochs, while spatial coexistence provides a direct way for several phenotypes to contribute jointly. However, neither temporal nor spatial organization maps uniquely onto a particular value of *m*, since temporally expressed phenotypes can also contribute jointly when they recur within the timescale over which new mutations experience selection. The same principle applies to environmentally induced phenotypes when different environmental conditions recur within that timescale. Intermediate values of *m* describe cases in which some, but not all, of the phenotype repertoire contributes jointly within that timescale. As *m* increases, the pleiotropic effects of a mutation across more task-specific phenotypes are exposed to selection within the same selective epoch.

The life cycle of *Plasmodium* illustrates how such an intermediate regime can arise. As discussed in the previous section, the repeatedly recurring ring, trophozoite, and schizont states may contribute jointly to selection during the asexual cycle, whereas sexual and mosquito-stage functions occur in other parts of the life cycle and are not realized during every asexual cycle [Howick et al., 2019]. The full life cycle can therefore include some stage-specific phenotypes whose fitness effects are repeatedly exposed to selection together and others whose effects are separated across longer timescales, corresponding in our model to 1 *< m < T* .

Other forms of life-cycle organization can likewise alter how cellular phenotypes are exposed to selection. In the choanoflagellate *Choanoeca flexa*, for example, reversible phenotypic plasticity allows colonies to switch rapidly between feeding and swimming conformations in response to environmental cues [Brunet et al., 2019, Reyes-Rivera et al., 2022]. The ichthyosporean *Chromosphaera perkinsii*, in contrast, undergoes transient multicellular development in which distinct cell types coexist within the same colony [Olivetta et al., 2024]. Such variation in the temporal and spatial organization of cellular phenotypes can change how much of the phenotype repertoire contributes to fitness within the same selective epoch. In our model, increasing the number of phenotypes that contribute jointly to fitness gradually relieves the constraint imposed by limited exposure to selection, allowing greater differentiation and optimization before the *m* = *T* limit is reached.

## Concluding remarks

The diversity of cellular states in unicellular organisms has motivated the hypothesis that spatial cell differentiation evolved through the integration of temporally differentiated ancestral states [Brunet and King, 2017, Mikhailov et al., 2009, Sebé-Pedrós et al., 2017, Zakhvatkin, 1949]. Our results suggest that such integration can change the constraints on the evolution of differentiation. Even when a shared genome contains enough programs to generate distinct cellular phenotypes, selection acting through those phenotypes separately can constrain their differentiation because the pleiotropic effects of mutations on other phenotypes are not exposed to selection at the same time. As more phenotypes contribute jointly to fitness, these effects are exposed to selection together, allowing greater differentiation under divergent functional demands until the number of available programs becomes limiting.

Temporal and spatial differentiation need not achieve this joint contribution through the same regulatory mechanisms. Rapid phenotypic switching in choanoflagellates can occur without de novo transcription [Brunet et al., 2021], while the feeding–swimming transition of *Choanoeca flexa* is controlled through sensory signaling and actomyosin contractility [Brunet et al., 2019, Reyes-Rivera et al., 2022]. Trypanosomes likewise regulate major developmental transitions between life-cycle stages predominantly post-transcriptionally, with protein-coding genes largely organized into polycistronic transcription units [Clayton, 2019, Kabani et al., 2009]. Stable animal cell types, in contrast, often rely on cell-type-specific transcriptional regulatory programs that maintain and individuate cell-type identity [Arendt et al., 2016]. Our model treats programs as reusable components of cellular phenotype without specifying how their deployment is regulated.

We impose *m*, the number of cellular phenotypes contributing jointly to fitness, for each simulation. The model therefore does not describe how the degree of joint contribution changes during a transition from temporally differentiated cell states to spatially differentiated cell types, or how regulation and genetic architecture evolve during such a transition. For example, mutations change the contributions of existing loci to programs but cannot duplicate loci, so the model does not describe how gene duplication and subsequent divergence create new opportunities for functional specialization [Arendt et al., 2016, Force et al., 1999]. Within this fixed genetic architecture, however, increasing the number of phenotypes that contribute jointly to fitness is sufficient to increase differentiation under divergent functional demands. Our results therefore suggest that the integration of ancestral cell states could have favored greater differentiation by allowing more of those phenotypes to contribute jointly to fitness, even without an increase in the number of available programs.

## Data availability

All results presented here are based on simulated data. The simulation and analysis code is publicly available at https://github.com/applied-phylo-lab/GEP_Evolutionand archived at Zenodo (https://doi.org/10.5281/zenodo.22782584, v1.0.0). The repository includes scripts that regenerate every simulated condition (make_data.py) and every figure (make_figures.py).

## Acknowledgments

We thank Rex Jiang, Joe Parker, Jaeda Patton, and members of the Pennell lab for critical feedback on this work and manuscript. The authors used ChatGPT and Claude to aid in searching the literature, editing the text for grammar and clarity, and developing the code. The authors take full responsibility for the content of the manuscript. This work was supported by a NIGMS award to MP (R35GM151348).

## 1 Simulation

The genotype–phenotype map and evolutionary dynamics are described in the main text under Genotype, regulation, and phenotype and Evolutionary dynamics and simulation.

### 1.1 Simulation parameters

Table S1 summarizes the parameters used in the main analyses and the values changed in the supplementary analyses.

### 1.2 Simulation length and termination

The maximum number of substitutions varied with *T* and *m* according to

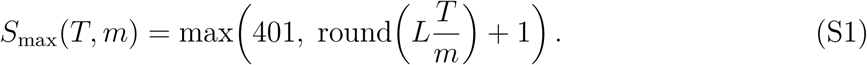

When only a small fraction of tasks contributes to fitness during each selective epoch, any particular task is encountered less frequently. We therefore allowed longer simulations at smaller *m*, with the maximum number of substitutions increasing in proportion to *T/m*. The substitution limit was at least 401 for every combination of *T* and *m*, unless the simulation terminated earlier, allowing comparisons at 50 and 400 substitutions across all conditions. For example, under sequential selection, the maximum is 401 substitutions for *T* = 2 and 4, 601 for *T* = 6, and 801 for *T* = 8. Under simultaneous selection, the maximum is 401 substitutions for every value of *T* .

Termination also accounted for changes in the active tasks. Under sequential and partially simultaneous selection, the absence of a beneficial mutation for the current task set does not imply that further adaptation is impossible. We therefore drew another set of active tasks whenever no beneficial mutation was available for the current set. Simulations ended when the maximum number of substitutions was reached or when no single-entry mutation was beneficial under any possible combination of *m* active tasks. We tested the latter condition exactly over all possible combinations of *m* active tasks. Simulations were also terminated after 100 consecutive task-set draws with no beneficial mutation. For populations that reached a local fitness peak, we retained their final phenotype values in subsequent fixed-substitution comparisons. For populations terminated after 100 consecutive unsuccessful task-set draws, we likewise retained their final recorded phenotype values in these comparisons.

### 1.3 Task ensembles and replicate populations

Task optima were generated from symmetric Dirichlet distributions as described under Task ensemble construction in the main text. For each combination of *T* and prescribed Δ*T*, the concentration parameter *α* was calibrated by bisection in log-space over *α* ∈ [10*−*2, 102]. At each step, 30 task ensembles were generated using the same set of random seeds, and the search continued until their average mean pairwise separation was within 0.02 of the prescribed value, with a maximum of 50 iterations.

Each replicate population was assigned an independently drawn task ensemble at the calibrated value of *α* and an independently drawn initial genotype. For comparisons among values of *m*, we used the same initial genotype and task ensemble for the corresponding population in each selection regime. The realized mean pairwise separation among task optima varied across replicate populations, even at the same value of *α*. To account for this variation, we normalized phenotypic differentiation in each population by the mean pairwise separation among its own task optima. Values of Δ*T* shown in the figures refer to the prescribed values.

### 1.4 Summary statistics

Both normalized differentiation and optimization range from zero to one. For a given genome, each phenotype **z***_j_*(*G*) is the Euclidean projection of its task optimum **T***_j_* onto the convex cone *{G***a** : **a** *≥* 0*}*. Projection onto this cone cannot increase pairwise distances, so

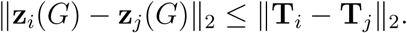

Averaging over task pairs therefore gives 0 ≤ Δ̄Z*̄*/Δ̄*T̄* ≤ 1.

For optimization, choosing all program coefficients to be zero is always possible, and each task optimum has unit length. The minimum distance to each optimum therefore satisfies

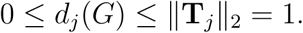

It follows that ‖*d*(*G*)‖_2_ ≤ √*T*, which gives 0 ≤ Ω ≤ 1.

In Figures 2–4 of the main text and Figures S2–S5, lines show means and shaded bands indicate 1 sample standard deviation across 200 replicate populations. The trajectory bands in Figure S1A–C instead show ±1 standard error of the mean, calculated as the sample standard deviation divided by the square root of the number of contributing populations at each substitution. For Figure S6, we calculated gains relative to sequential selection by subtracting each replicate population’s value at *m* = 1 from its value at each *m*. Corresponding populations shared the same initial genotype and task ensemble across selection regimes. Lines show the mean of these paired differences across 200 populations, and shaded bands indicate ±1 standard error, calculated as the sample standard deviation of the paired differences divided by √200. The gain is zero at *m* = 1 by construction.

### 1.5 Evolutionary trajectories and comparison points

To set comparison points for earlier and later stages of adaptation, we examined how differentiation, optimization, and the fraction of beneficial mutations changed over successive substitutions (Figure S1). For each substitution, we calculated the fraction of the *LK* possible single-entry mutations that would increase fitness under the active task set. This fraction was recorded for the task set that produced the substitution (Figure S1C); draws with no beneficial mutation were not included. Under simultaneous selection, populations reached local fitness peaks before 400 substitutions, after which the phenotypic measures no longer changed (Figure S1A–C, see solid lines). Under sequential and intermediate selection, the fraction of beneficial mutations declined rapidly early in adaptation and changed more slowly later, while remaining above zero (Figure S1C, see dotted and dashed lines). The phenotypic measures showed a similar pattern (Figure S1A–B), although differentiation continued to increase under some conditions (Figure S1A). We therefore used 50 and 400 substitutions as heuristic comparison points for earlier and later stages of adaptation, without assuming that all trajectories had strictly stabilized by the later point.

---

**Figure S1.**
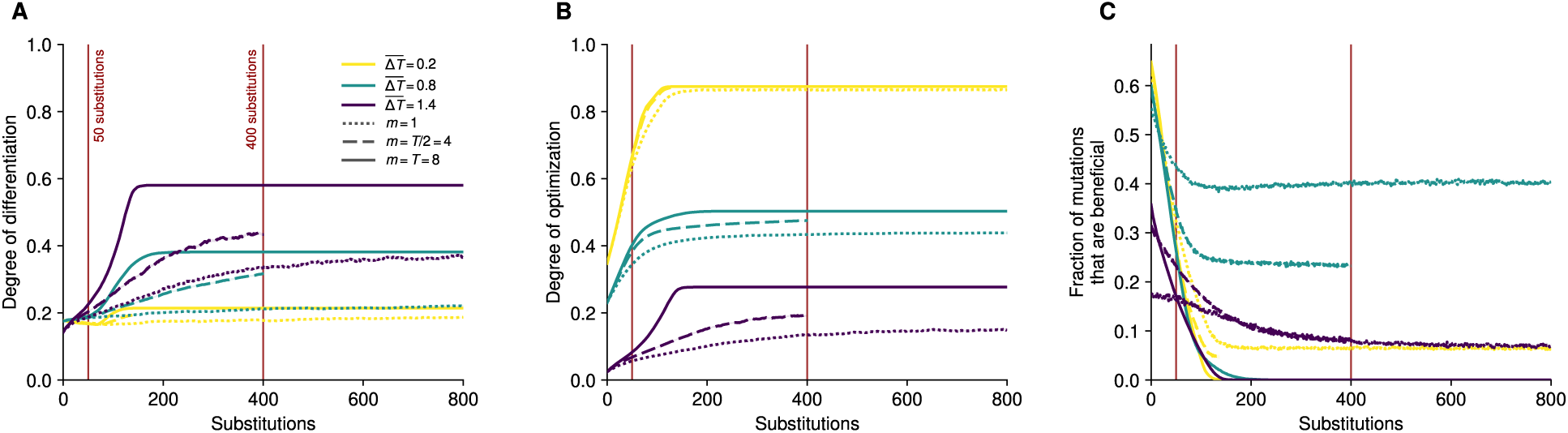
Evolutionary trajectories and the choice of comparison points. **A–C.** Phenotypic differentiation, optimization, and the fraction of beneficial single-entry mutations over successive substitutions at *T* = 8, with *K* = 4 and *ρ* = 0.25. Colors indicate the prescribed mean pairwise separation among task optima. Dotted, dashed, and solid lines show sequential selection (*m* = 1), selection on half the tasks (*m* = 4), and simultaneous selection (*m* = 8), respectively. Red vertical lines mark the 50- and 400-substitution comparisons. Lines show means and shaded bands indicate ±1 standard error across contributing populations, from 200 initial replicates per condition. In **C**, the fraction is the number of single-entry mutations that increase fitness under the active task set divided by *LK* = 400. It is recorded for the task set preceding each substitution, excluding task-set draws with no beneficial mutation. Under simultaneous selection, populations reach local fitness peaks where no single-entry mutation can improve fitness. Their final phenotype values are retained thereafter, and their fraction of beneficial mutations is zero. The solid trajectories therefore become flat once all populations have reached these peaks; the flat portions do not represent further substitutions. The *m* = 4 simulations were run for at most 401 substitutions, so the dashed trajectories at the two stronger target separations end near 400. At the weakest separation, the dashed trajectory ends earlier. Trajectories end when fewer than 90% of the original replicates remain available. **Alt text:** Three plots show phenotypic differentiation, optimization, and the fraction of beneficial single-entry mutations against the number of substitutions, from zero to eight hundred. Colors indicate separation among task optima, from yellow for weakly separated optima to purple for strongly separated optima. Dotted, dashed, and solid lines show sequential selection, intermediate selection on half the tasks, and simultaneous selection, respectively. Two dark red vertical lines mark the 50- and 400-substitution comparison points, representing the early and late phases of adaptation. In the first two plots, the solid curves rise steeply and then remain flat from roughly 150 substitutions onward, while the dotted and dashed curves continue to change more slowly. In the third plot, the fraction of beneficial single-entry mutations falls steeply from its initial value, reaching zero under simultaneous selection and settling at low positive values under sequential and intermediate selection. Shading shows the standard error across contributing populations.

**Figure S2.**
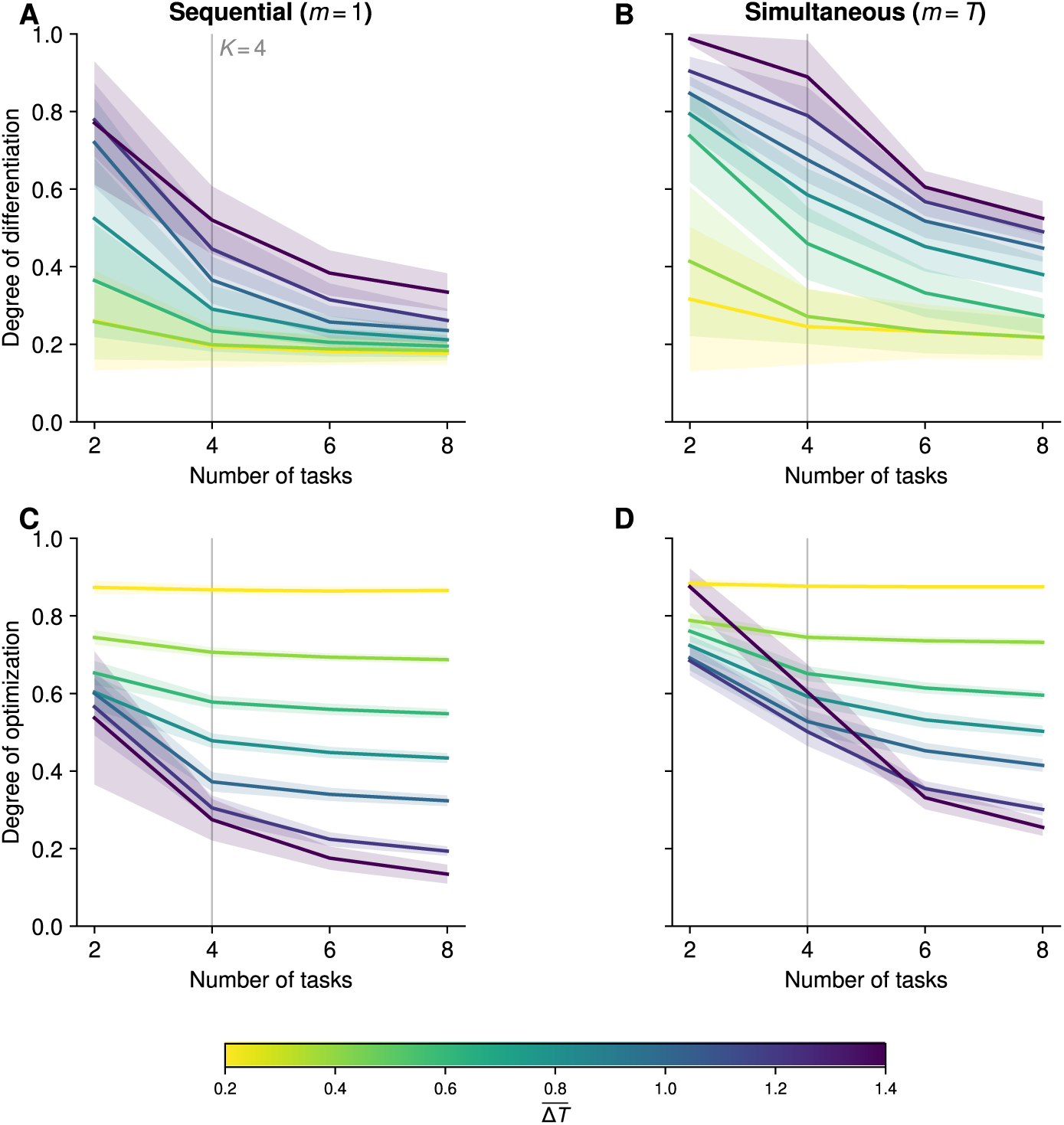
Sequential and simultaneous selection under a negative power-mean fitness function. Fitness is calculated using a power mean with *r* = −2, which gives greater weight to poorly performing tasks. **A, B.** Phenotypic differentiation, Δ̄Z*̄*/Δ̄*T̄*, after 400 substitutions under sequential (**A**) and simultaneous (**B**) selection. **C, D.** Optimization index, Ω, under the same conditions. Colors indicate target separation, Δ̄*T̄*. Vertical lines mark *T* = *K* = 4. Under sequential selection, fitness depends on a single task and is therefore independent of *r*. Panels **A** and **C** repeat Figure 2C,E of the main text for comparison. Lines show means and shaded bands indicate ±1 standard deviation across 200 replicate populations. Other parameters are unchanged from Figure 2 (*L* = 100, *K* = 4, *γ* = 1, *ρ* = 0.25, *µ* = 10*−*7, and *N* = 104). **Alt text:** Four plots compare sequential selection on the left with simultaneous selection on the right, showing differentiation in the top row and optimization in the bottom row against task number. Colors run from yellow for weakly separated task optima to purple for strongly separated optima. Grey vertical lines mark four tasks, equal to the number of programs. Under sequential selection, differentiation falls steeply as tasks accumulate, and the differences among separations narrow. Under simultaneous selection, differentiation remains near its maximum at strongly separated optima while tasks do not outnumber programs, and declines thereafter. Optimization is high and nearly flat for weakly separated optima in both regimes, and declines with task number when optima are strongly separated, most steeply under simultaneous selection. Shading shows one standard deviation across replicate populations.

**Figure S3.**
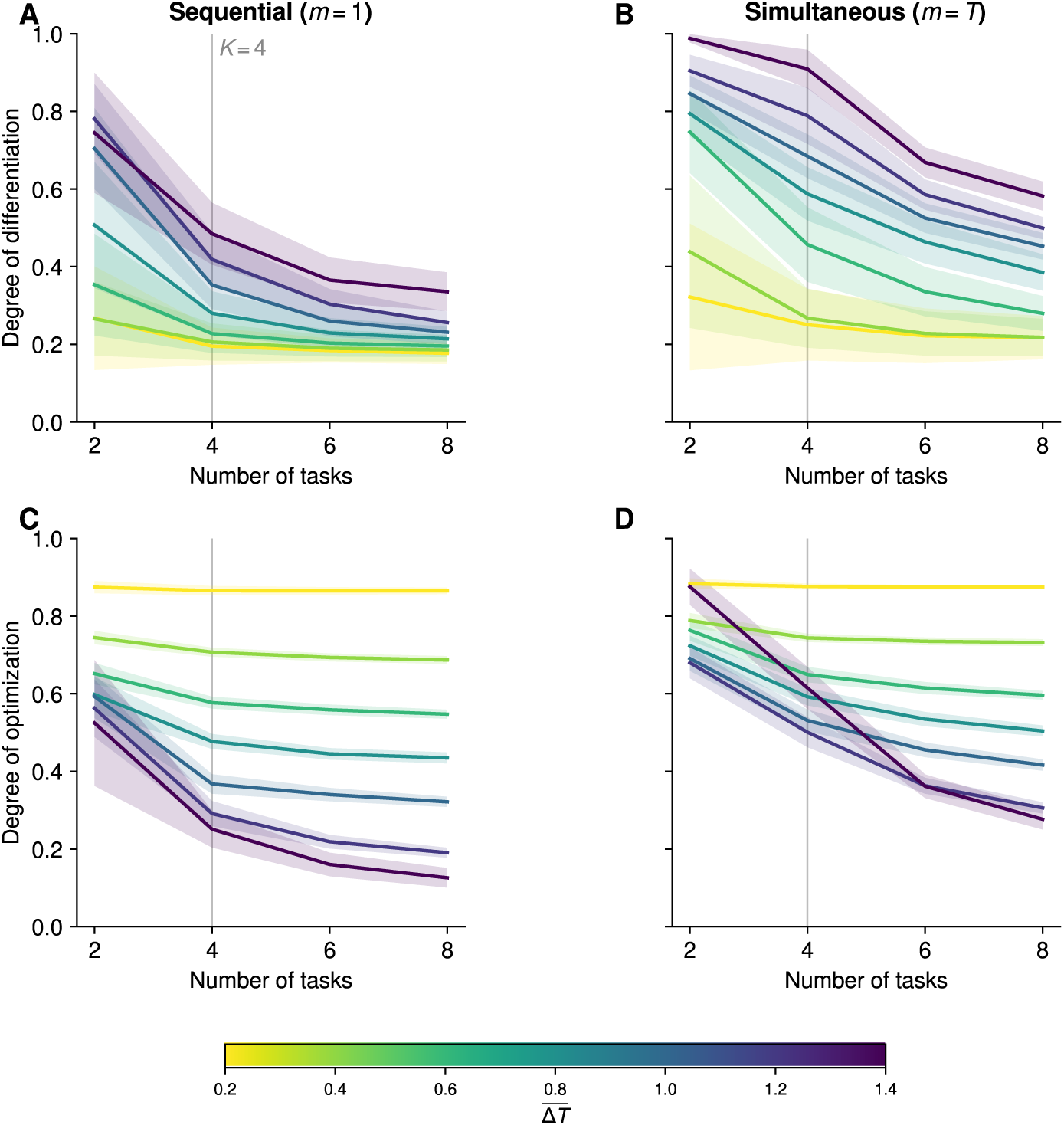
Sequential and simultaneous selection with a more strongly curved performance function. Task performance is calculated using *γ* = 4, so performance declines more sharply with distance from the task optimum. Fitness is the geometric mean of performance across active tasks (*r* = 0). **A, B.** Phenotypic differentiation, Δ̄Z*̄*/Δ̄*T̄*, after 400 substitutions under sequential (**A**) and simultaneous (**B**) selection. C**, D.** Optimization index, Ω, under the same conditions. Colors indicate target separation, *Δ̄*T̄**. Vertical lines mark *T* = *K* = 4. Lines show means and shaded bands indicate ±1 standard deviation across 200 replicate populations. Other parameters are as in Figure 2 of the main text. **Alt text:** Four plots compare sequential selection on the left with simultaneous selection on the right, showing differentiation in the top row and optimization in the bottom row against task number. Colors run from yellow for weakly separated task optima to purple for strongly separated optima. Grey vertical lines mark four tasks, equal to the number of programs. With a more sharply declining performance function, the contrast between the two regimes is unchanged. Under sequential selection, differentiation falls steeply as tasks accumulate, and the differences among separations narrow. Under simultaneous selection, differentiation remains near its maximum at strongly separated optima while tasks do not outnumber programs, and declines thereafter. Optimization is high and nearly flat for weakly separated optima and declines with task number for strongly separated optima, most steeply under simultaneous selection. Shading shows one standard deviation across replicate populations.

**Figure S4.**
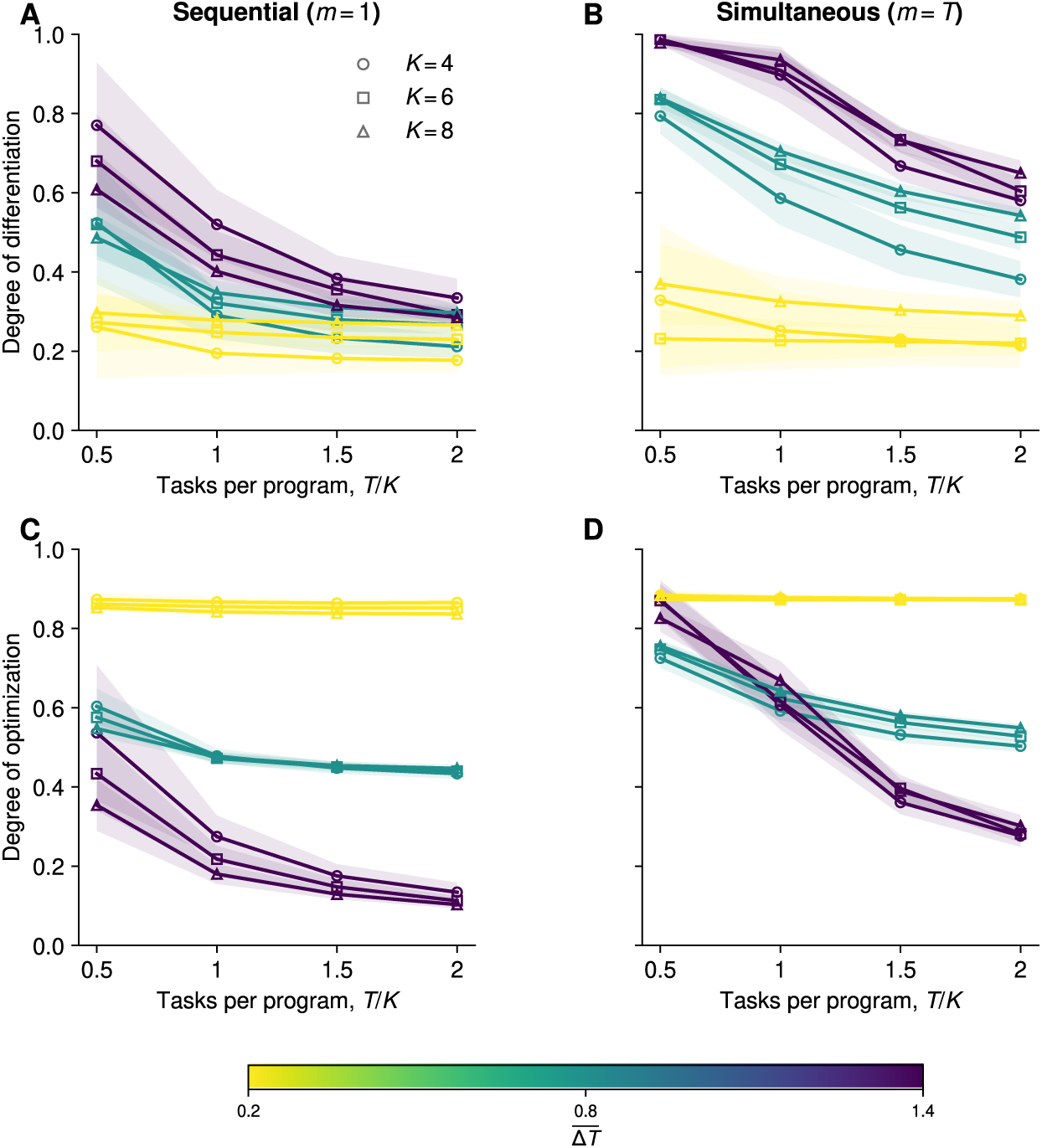
Phenotypic evolution with different numbers of available programs. Sequential and simultaneous selection are compared for *K* = 4, 6, and 8 after 400 substitutions. **A, B.** Phenotypic differentiation, ΔZ*̄*/Δ*T̄*, under sequential (**A**) and simultaneous (**B**) selection. **C, D.** Optimization index, Ω, under the same conditions. Task number is expressed relative to program number, *T/K*, with equal numbers of tasks and programs at *T/K* = 1. Circles, squares, and triangles indicate *K* = 4, 6, and 8, respectively. Colors indicate target separation, Δ*T̄* = 0.2, 0.8, and 1.4, as in Figures 2 and 3 of the main text. The initial density is *ρ* = 0.25 for all values of *K*. Lines show means and shaded bands indicate ±1 standard deviation across 200 replicate populations. Other parameters are as in Figure 2. **Alt text:** Four plots compare sequential selection on the left with simultaneous selection on the right, showing differentiation in the top row and optimization in the bottom row against tasks per program, from one half to two. Circles, squares, and triangles distinguish four, six, and eight programs. Colors indicate weak, intermediate, and strong separation among task optima. Equal numbers of tasks and programs fall at one task per program. Under sequential selection, differentiation declines with increasing tasks per program at every program number. Under simultaneous selection, differentiation remains near its maximum at strongly separated optima up to one task per program, and trajectories for the three program numbers lie close to one another when plotted against tasks per program. Optimization declines with increasing tasks per program in both regimes, most steeply for strongly separated optima under simultaneous selection. Shading shows one standard deviation across replicate populations.

**Figure S5.**
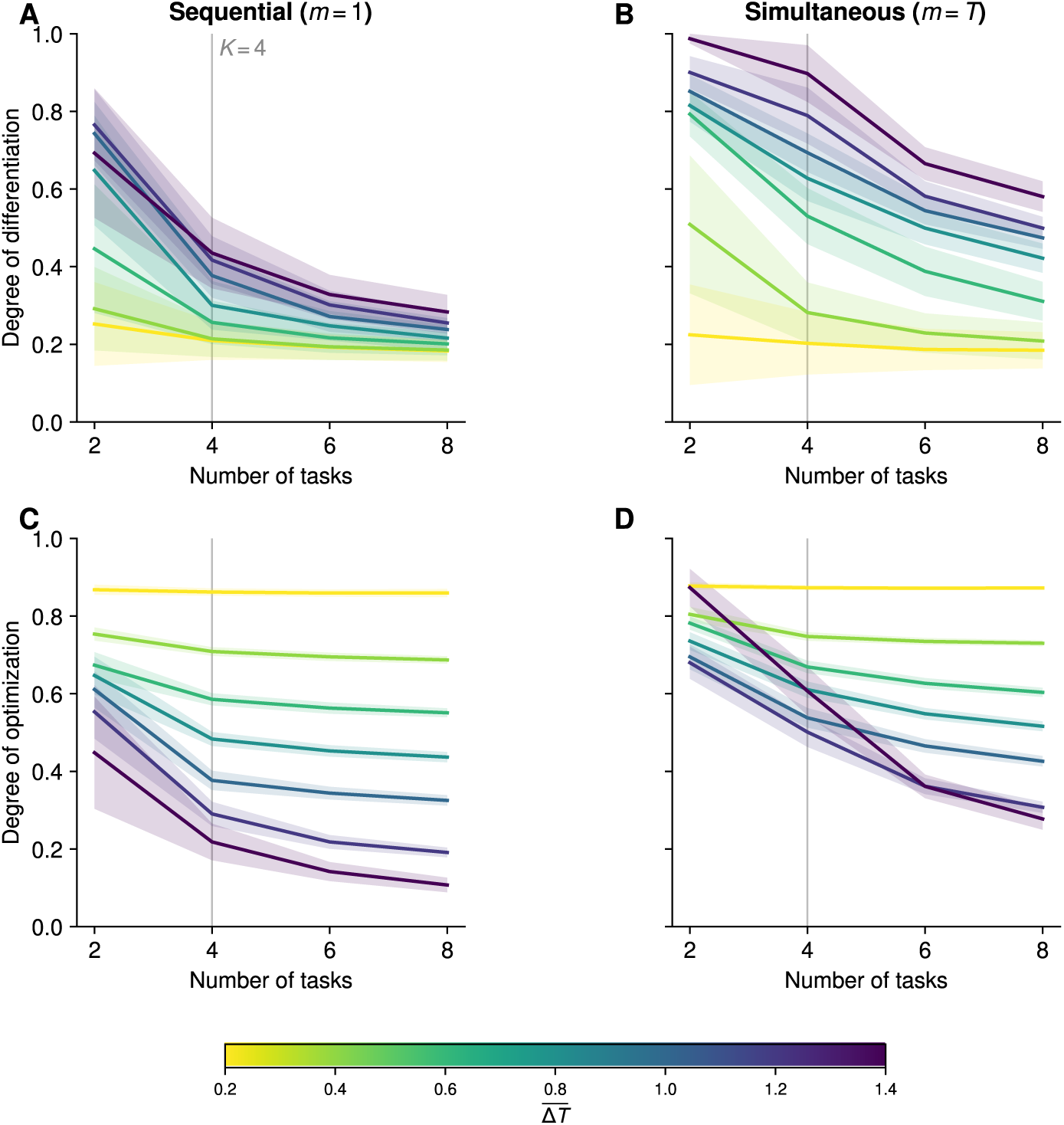
Phenotypic evolution from a denser initial genotype matrix. The initial density is *ρ* = 0.5, so each locus affects two programs on average at *K* = 4, compared with one under the *ρ* = 0.25 initialization. **A, B.** Phenotypic differentiation, ΔZ*̄*/Δ*T̄*, after 400 substitutions under sequential (**A**) and simultaneous (**B**) selection. **C, D.** Optimization index, Ω, under the same conditions. Colors indicate target separation, Δ*T̄*. Vertical lines mark *T* = *K* = 4. Lines show means and shaded bands indicate ±1 standard deviation across 200 replicate populations. Other parameters are as in Figure 2 of the main text. **Alt text:** Four plots compare sequential selection on the left with simultaneous selection on the right, showing differentiation in the top row and optimization in the bottom row against task number. Colors run from yellow for weakly separated task optima to purple for strongly separated optima. Grey vertical lines mark four tasks, equal to the number of programs. Starting from a denser genotype matrix does not change the contrast between the regimes. Under sequential selection, differentiation falls steeply as tasks accumulate, and the differences among separations narrow. Under simultaneous selection, differentiation remains near its maximum at strongly separated optima while tasks do not outnumber programs, and declines thereafter. Optimization is high and nearly flat for weakly separated optima and declines with task number for strongly separated optima, most steeply under simultaneous selection. Shading shows one standard deviation across replicate populations.

**Figure S6.**
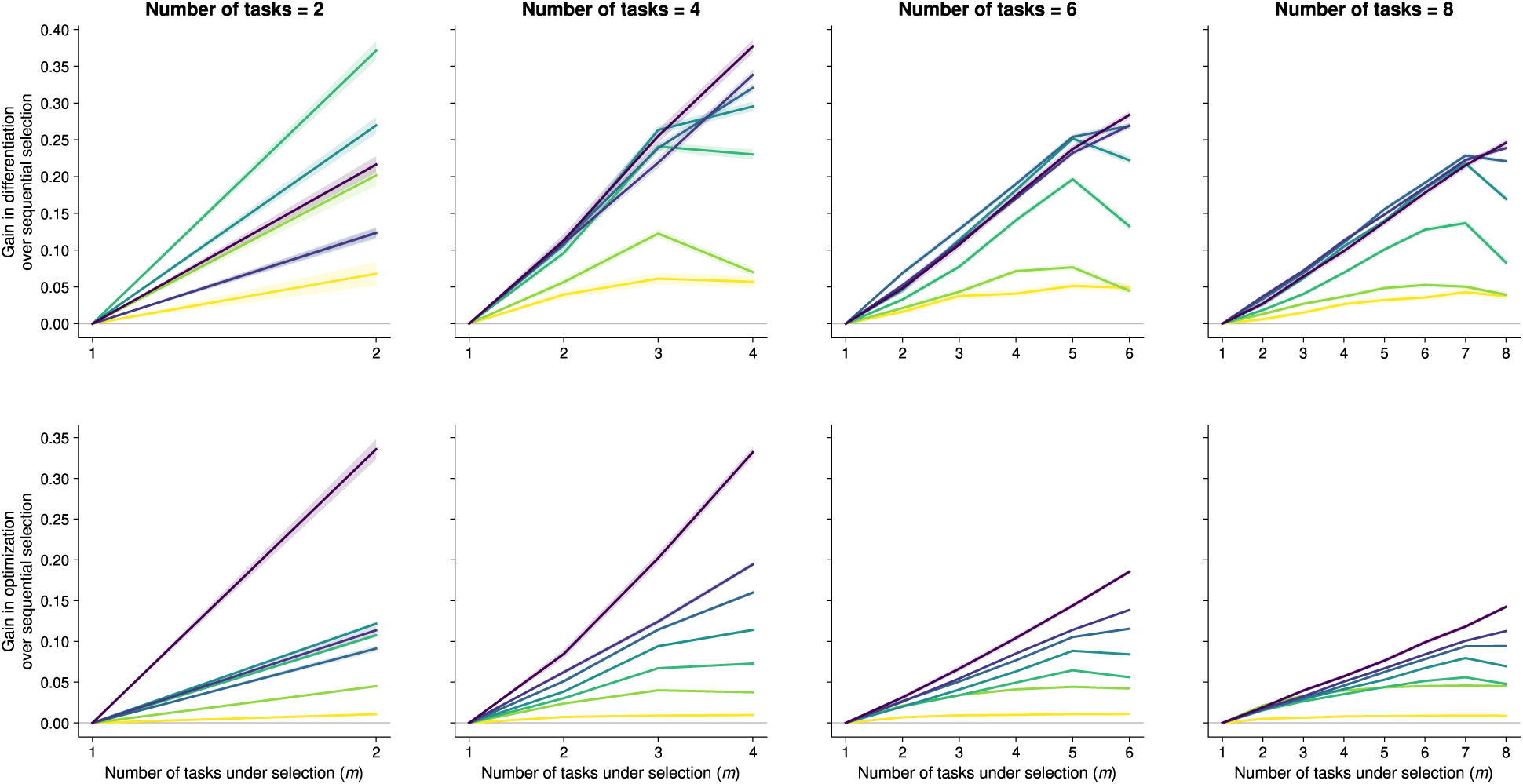
Gains in differentiation and optimization as more phenotypes contribute jointly to fitness. Gains relative to sequential selection are shown against *m*, the number of task-specific phenotypes contributing jointly to fitness, after 400 substitutions. The top row shows the gain in phenotypic differentiation, ΔZ*̄*/Δ*T̄*, and the bottom row shows the gain in optimization, Ω. Columns indicate task number, *T*, with *K* = 4 throughout. Colors indicate target separation, Δ*T̄*. For each replicate, the gain is calculated by subtracting the value at *m* = 1 from that at each *m*. Gains are therefore zero at *m* = 1, and *m* = *T* corresponds to simultaneous selection. At strong target separation, gains are greatest at *m* = *T* ; at weaker separation, they peak at intermediate *m*. Replicates share the same initial genotype and task ensemble across values of *m*. Lines show mean paired differences and shaded bands indicate ±1 standard error across 200 replicate populations. Simulations are the same as those used in Figure 4 of the main text. Parameters are as in Figure 2 and colors follow Figure 2’s scale. **Alt text:** Eight plots show the gain over sequential selection, with differentiation in the top row and optimization in the bottom row, against the number of task-specific phenotypes contributing jointly to fitness. Columns correspond to two, four, six, and eight tasks. Colors run from yellow for weakly separated task optima to purple for strongly separated optima. Every curve starts at zero, since the gain is measured relative to selection on one phenotype at a time. From four tasks onward, gains in both measures are larger at more strongly separated optima. At strongly separated optima, the gains increase as more phenotypes contribute jointly and are greatest when all of them do. At weaker separations, the curves instead peak before all phenotypes contribute jointly and then decline, an effect that is more pronounced for differentiation than for optimization. Shading shows the standard error of the paired differences across replicate populations.

**Table S1.** Parameters used in the main and supplementary analyses.

| Symbol | Meaning | Main | Supplementary |
| --- | --- | --- | --- |
| $L$ | Number of genetic loci | 100 | — |
| $K$ | Number of programs | 4 | 6, 8 |
| $T$ | Number of tasks | 2, 4, 6, 8 | 3, 6, 9, 12 at $K = 6$;<br>4, 8, 12, 16 at $K = 8$ |
| $m$ | Number of task-specific phenotypes contributing jointly to fitness | 1 to $T$ | $m \in \{1, T\}$ for supplementary robustness analyses |
| $\overline{\Delta T}$ | Mean pairwise separation among task optima | 0.2–1.4 in increments of 0.2 | same |
| $\rho$ | Initial density of the genotype matrix | 0.25 | 0.5 |
| $\gamma$ | Performance-function parameter | 1 | 4 |
| $r$ | Power-mean parameter | 0 | −2 |
| $N$ | Effective population size | $10^4$ | — |
| $\mu$ | Per-site per-generation mutation rate | $10^{-7}$ | — |
| $n_{\text{rep}}$ | Number of replicate populations per condition | 200 | — |

